# The *MUC5B* promoter variant results in proteomic changes in the non-fibrotic lung

**DOI:** 10.1101/2024.11.26.625453

**Authors:** Jeremy A. Herrera, Mark Maslanka, Rachel Z. Blumhagen, Rachel Blomberg, Nyan Ye Lwin, Janna Brancato, Carlyne D. Cool, Jonathan P. Huber, Jonathan S. Kurche, Chelsea M. Magin, Kirk C. Hansen, Ivana V. Yang, David A. Schwartz

## Abstract

The gain-of-function *MUC5B* promoter variant is the dominant risk factor for the development of idiopathic pulmonary fibrosis (IPF). However, its impact on protein expression in both non-fibrotic control and IPF lung specimens have not been well characterized. Utilizing laser capture microdissection coupled to mass spectrometry (LCM-MS), we investigated the proteomic profiles of airway and alveolar epithelium in non-fibrotic controls (n = 12) and IPF specimens (n = 12), stratified by the presence of the *MUC5B* promoter variant. Through qualitative and quantitative analyses, as well as pathway analysis and immunohistological validation, we have identified a distinct MUC5B-associated protein profile. Notably, the non-fibrotic control alveoli exhibited substantial MUC5B-associated protein changes, with an increase of IL-3 signaling. Additionally, we found that the epithelial cells overlying IPF fibroblastic foci cluster closely to alveolar epithelia and express proteins associated with cellular stress pathways. In conclusion, our findings suggest that the *MUC5B* promoter variant leads to protein changes in alveolar and airway epithelium that appears to be associated with the initiation and progression of lung fibrosis.

## Introduction

Usual interstitial pneumonia (UIP) is a broad category of fibrotic lung diseases that have a poor clinical outcome. IPF, the most severe form of lung fibrosis, is defined radiographically and pathologically as UIP [1]. Although IPF is traditionally considered idiopathic, emerging literature indicates that genetic factors account for at least 30% of the risk of developing IPF [2–5], and the *MUC5B* promoter variant is responsible for approximately 50% of the genetic risk of IPF [3]. Thus, understanding the spatial lung proteome in the context of the *MUC5B* promoter variant should help us decipher critical elements of protein biology in IPF.

We have previously developed laser capture microdissection coupled mass spectrometry (LCM-MS) for formalin-fixed and -stained lung tissue [6]. This method allowed us to determine the proteomic profiles of the two lesions of UIP/IPF, the honeycomb cyst [7] and fibroblastic foci [8]. Considering the potential role of the *MUC5B* promoter variant in IPF pathogenesis, we investigated proteomic differences in characteristic IPF lesions in relation to the *MUC5B* promoter variant. In addition, we performed LCM-MS to analyze the aberrant basaloid epithelial cells/transitional cells that overlie fibroblastic foci [9–12]; herein we refer to these as epithelia overlying fibroblastic foci. Our specimens were balanced for the *MUC5B* promoter variant in non-fibrotic control and IPF samples.

## Results

### The *MUC5B* promoter variant is associated with protein changes in non-fibrotic control lungs

We performed LCM-MS analysis on small airways and alveoli from non-fibrotic control specimens (n = 12 balanced for the *MUC5B* promoter variant) to determine which proteins are associated with the *MUC5B* promoter variant. Of these specimens, 6 where homozygous for the wildtype allele (GG) and 6 heterozygous for the MUC5B promoter variant (GT). We found that two proteins are decreased in non-fibrotic control small airways with the *MUC5B* promoter variant compared to without: small ribosomal subunit protein eS4, Y isoform 1 (RPS4Y1) and protein canopy homolog 2 (CNPY2) (**Figure 1A**; a full list of differentially expressed proteins for all *MUC5B* promoter variant comparisons are found in **Supplemental Table 1**).

**Figure 1:**
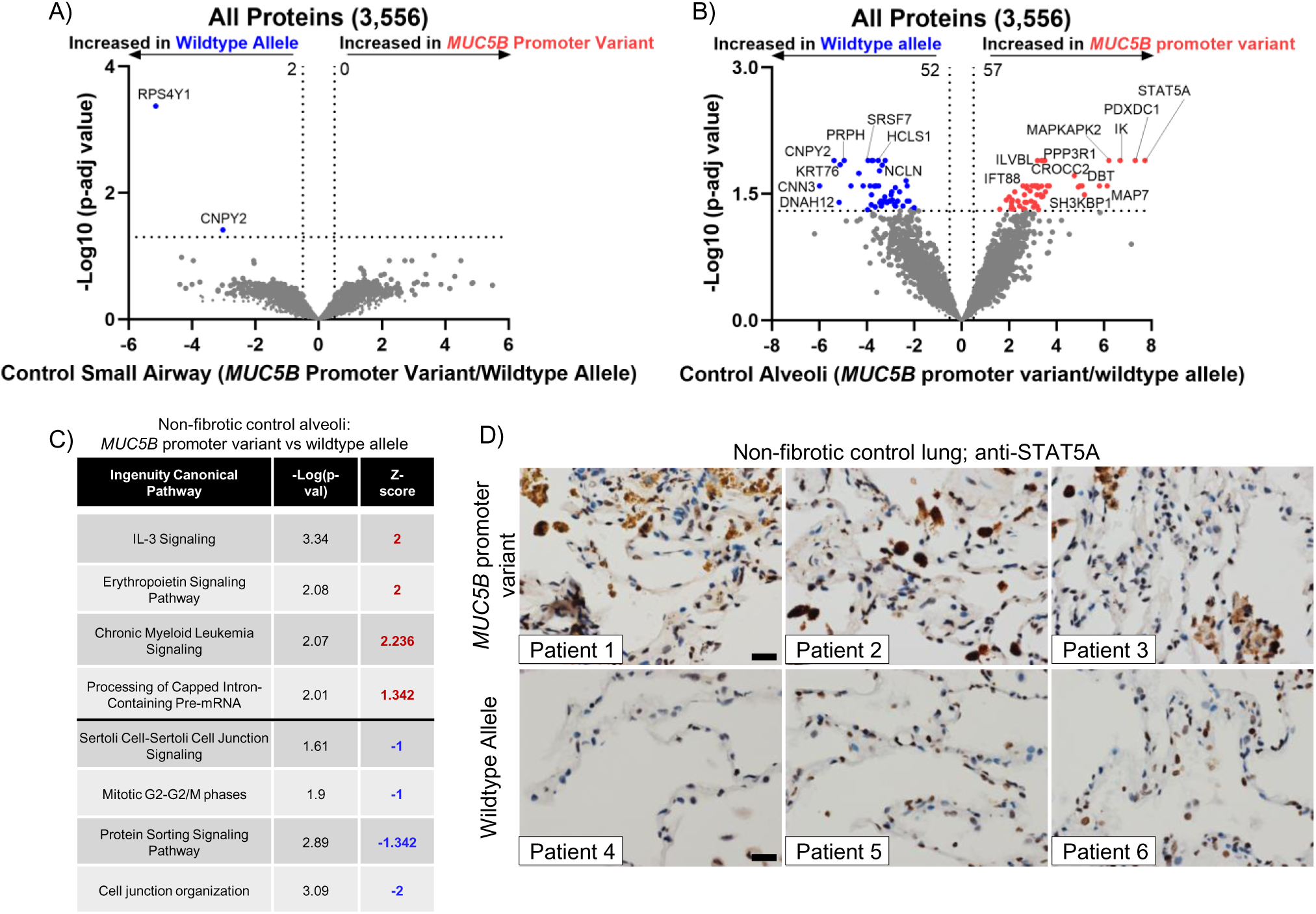
Impact of the *MUC5B* promoter variant on non-fibrotic control lung: Volcano plot comparing the *MUC5B* promoter variant and wildtype allele in non-fibrotic control (**A**) small airways and (**B**) alveoli, showing the negative natural log of p-adjusted values (0.05) for each protein. (**C**) Ingenuity pathway analysis showing the top 4 most upregulated (positive Z-score in red font) or top 4 most downregulated (negative Z-score in blue font) pathways in non-fibrotic control alveoli. N = 12 specimens (6 wildtype and 6 *MUC5B* promoter variant). (**D**) Immunohistochemistry against STAT5A on non-fibrotic control lungs (n = 6 specimens, balanced for the *MUC5B* promoter variant). Scale bar represents 20 microns.

In addition, we report that the *MUC5B* promoter variant is associated with 109 protein changes in non-fibrotic control alveoli (**Figure 1B**). The most significantly increased protein is a signal transducer and activator of transcription 5A (STAT5A) which functions as a transcription factor. Also increased by the *MUC5B* promoter variant is mitogen-activated protein kinase–activated protein kinase–2 (MAPKAPK2). MAPKAPK2 has been shown to be phosphorylated in IPF epithelial cells and its inhibition reduces bleomycin-induced lung injury [13]. Pathway analysis via Ingenuity Pathway Analysis (IPA) demonstrate that ‘IL-3 signaling’ is the most significantly increased pathway, whereas ‘cell junction organization’ is the most decreased pathway in non-fibrotic control alveoli harboring the *MUC5B* promoter variant (**Figure 1C**). At the level of proteomics, MUC5B was not significantly changed in non-fibrotic control alveoli. We used immunohistochemistry (IHC) to validate our findings and show marked expression of STAT5A in non-fibrotic control lungs, particularly in immune cells, harboring the *MUC5B* promoter variant (**Figure 1D**). These protein changes in non-fibrotic lungs suggest that the *MUC5B* promoter variant may establish a vulnerable distal lung epithelium.

### The histopathological lesions of IPF uniquely cluster

To guide our understanding of IPF at the protein level, we performed hierarchical clustering of our regions of interest (IPF honeycomb cyst, IPF small airways, IPF fibroblastic foci, IPF alveoli, IPF epithelia overlying fibroblastic foci, non-fibrotic control small airway and non-fibrotic control alveoli) based on the top 1,000 variable proteins (**Figure 2**). These regions were histologically confirmed by a pathologist (representative laser captured images in **Supplementary Figure 1**). We found that these regions of interest cluster into 3 groups, with some deviations. Firstly, we found that IPF honeycomb cyst cluster with both IPF and non-fibrotic control small airways. We additionally found that the IPF fibroblastic foci samples cluster together. Lastly, we found that the alveolar samples cluster with the IPF epithelia overlying fibroblastic foci.

**Figure 2:**
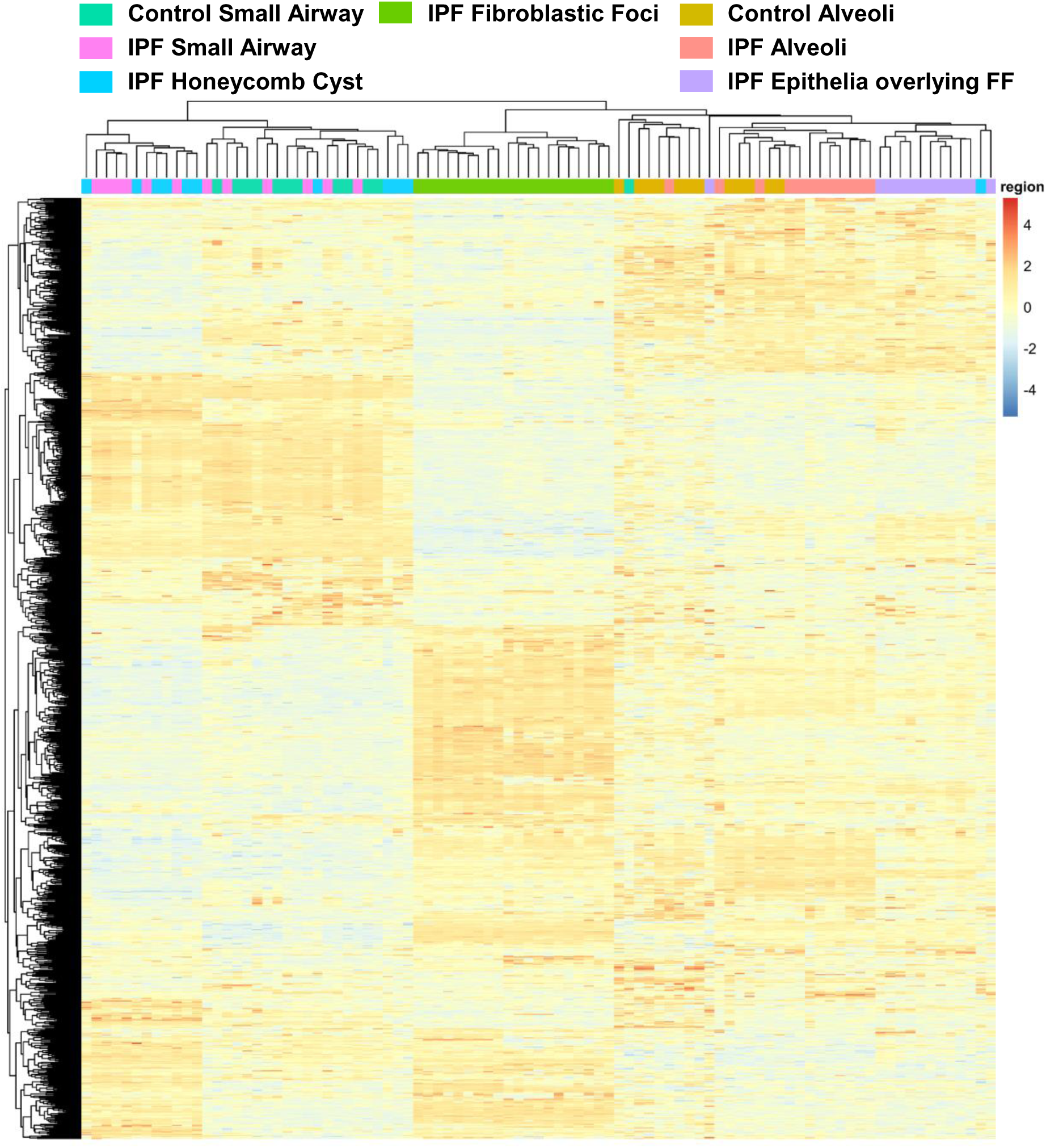
The histopathological lesions of IPF uniquely cluster. A hierarchical cluster analysis based on the top 1,000 variable proteins. For IPF specimens (n = 12), we collected 12 small airway, 11 honeycomb cyst, 12 epithelia overlying fibroblastic foci, 12 alveoli, and 12 fibroblastic foci samples from the same 12 IPF specimens. We further collected an additional 8 fibroblastic foci samples (a total of 20 fibroblastic foci samples) from 8 additional IPF specimens. For non-fibrotic control specimens (n = 12), we collected 12 small airway and 12 alveoli samples from the same specimen. The samples were balanced for the *MUC5B* promoter variant.

### Normal-appearing IPF epithelia (small airway and alveoli) are abnormal at the protein level

To understand the protein profiles of IPF airways, we first used a qualitative approach to determine which proteins are present in the airways. We considered a protein present if it was detected in 80% or more of samples within each group. Given this approach, we detected a total of 2,719 airway proteins when grouping IPF honeycomb cyst, IPF small airways, and non-fibrotic control small airways (**Supplemental Table 2**). Of the 2,719 total airway proteins, 175 were uniquely expressed in the normal-appearing IPF small airways (**Supplemental Figure 2A**). Gene enrichment analysis of the unique IPF small airway proteins showed that the most significantly upregulated pathways are ‘Anchoring of the basal body to the plasma membrane’ and ‘Cilium assembly’, which are pathways involved in ciliogenesis (**Supplemental Table 3**).

We next performed a quantitative analysis and found 124 significantly increased proteins in IPF small airways and 70 proteins increased in non-fibrotic control small airways (**Figure 3A**; a full list of differentially expressed proteins for all regions are found in **Supplemental Table 4**). The most significantly increased protein in IPF small airways is cilia-and flagella-associated protein 46 (CFAP46). Axin interactor, dorsilization-associated protein (AIDA) which has been shown to antagonize the JNK signaling pathway [14], is the most significantly decreased protein. IPF small airway pathway analysis demonstrated an increase of ‘Rho GTPases activate IQ motif containing GTPase activating proteins (IQGAPs)’ and ‘post-translational protein phosphorylation’ (**Figure 3B**). IQGAPs regulate many cellular processes including MAPK signaling pathways [15]. These results support the concept that the normal-appearing small airways in IPF are functionally abnormal, demonstrating elevated levels of components involved in signaling pathways and ciliogenesis. However, none of the 194 significantly changed proteins in IPF small airways were differentially regulated by the *MUC5B* promoter variant (**Figure 3C**).

**Figure 3:**
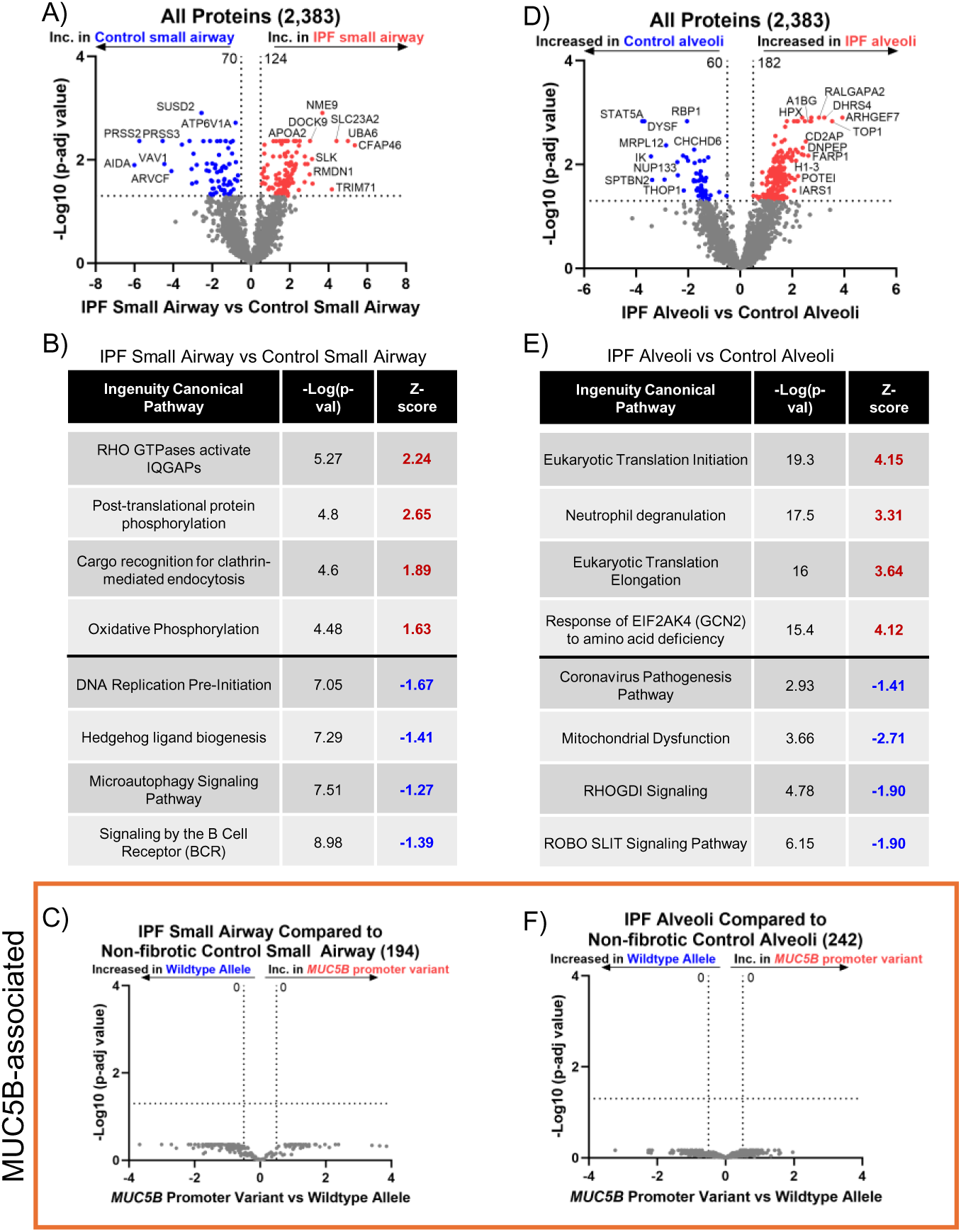
The normal epithelium (airway and alveoli) of IPF are abnormal. (**A & D**) Volcano plot comparing (A) IPF small airway and non-fibrotic control small airways, (D) IPF alveoli and non-fibrotic control alveoli, showing the negative natural log of p-adjusted values plotted against the base 2 log (fold change) for each protein. (**B & E**) Ingenuity pathway analysis showing the top 4 most upregulated (positive Z-score in red font) or top 4 most downregulated (negative Z-score in blue font) pathways per comparison. (**C & F**) Focusing on the significantly changed proteins from (A & D), we re-analyzed the data for the *MUC5B* promoter variant and show volcano plots for (C) IPF small airways and (F) IPF alveoli. N = 12 per group (6 wildtype and 6 *MUC5B* promoter variant).

To understand the protein profiles of the normal-appearing alveoli in IPF, we performed a qualitative analysis comparing the epithelia overlying fibroblastic foci with IPF and non-fibrotic control alveoli and detected a total of 1,996 proteins (**Supplemental Figure 2B**). This comparison is based on the observation that the epithelia overlying the fibroblastic foci are positive for alveolar markers [8, 16], and cluster with non-fibrotic control and IPF alveoli based on unsupervised analysis (**Figure 2**). We then performed a gene enrichment analysis of the 104 unique IPF alveolar proteins and found that ‘Regulation of insulin secretion’ and ‘Integration of energy metabolism’ are among the strongest enriched pathways.

To understand IPF alveoli in further detail, we quantitatively compared IPF alveoli to non-fibrotic control alveoli and found 242 differentially expressed proteins (**Figure 3D**). The most significantly increased protein in IPF alveoli is Rho guanine nucleotide exchange factor 7 (ARHGEF7) which is a guanine exchange factor for Rac1 and Cdc42 [17]; Cdc42 knockout in alveolar type II cells (AT2) drives periphery-to-center lung fibrosis [18]. Pathway analysis of IPF alveoli demonstrated an increase in a variety of pathways involved in translational control (e.g., eukaryotic translation initiation and elongation) (**Figure 3E**). Our results confirm that the normal-appearing IPF alveoli adjacent to fibroblastic foci are biologically abnormal with increased proteins involved in translational control and metabolism. However, none of the 242 differentially expressed alveolar proteins are regulated by the *MUC5B* promoter variant (**Figure 3F**).

### Membrane trafficking and remodeling of epithelial adherens junctions define IPF honeycomb cysts

We next focused our analysis on the IPF characteristic lesions: honeycomb cysts and fibroblastic foci (FF). To understand the biology of IPF honeycomb cysts, we first performed a gene enrichment analysis of the 258 uniquely expressed IPF honeycomb cyst proteins (**Supplemental Figure 2A**). The most significantly enriched pathway is ‘membrane trafficking’, a secretory membrane system that may reflect increased mucus production in this region. We next performed a quantitative analysis and found 187 significantly regulated proteins in IPF honeycomb cysts when compared to adjacent IPF small airways (**Figure 4A**). Several mucin or secretory-associated proteins are significantly increased in IPF honeycomb cysts, including MUC5B, MUC1, BPIFB1, SCGB3A1, and NAPSA. Pathway analysis of IPF honeycomb cyst (as compared to IPF small airway) showed enrichment of ‘remodeling of epithelial adherens junctions’ as the most significantly increased pathway whereas the most decreased pathway is ‘cilium assembly’ (**Figure 4B**).

**Figure 4:**
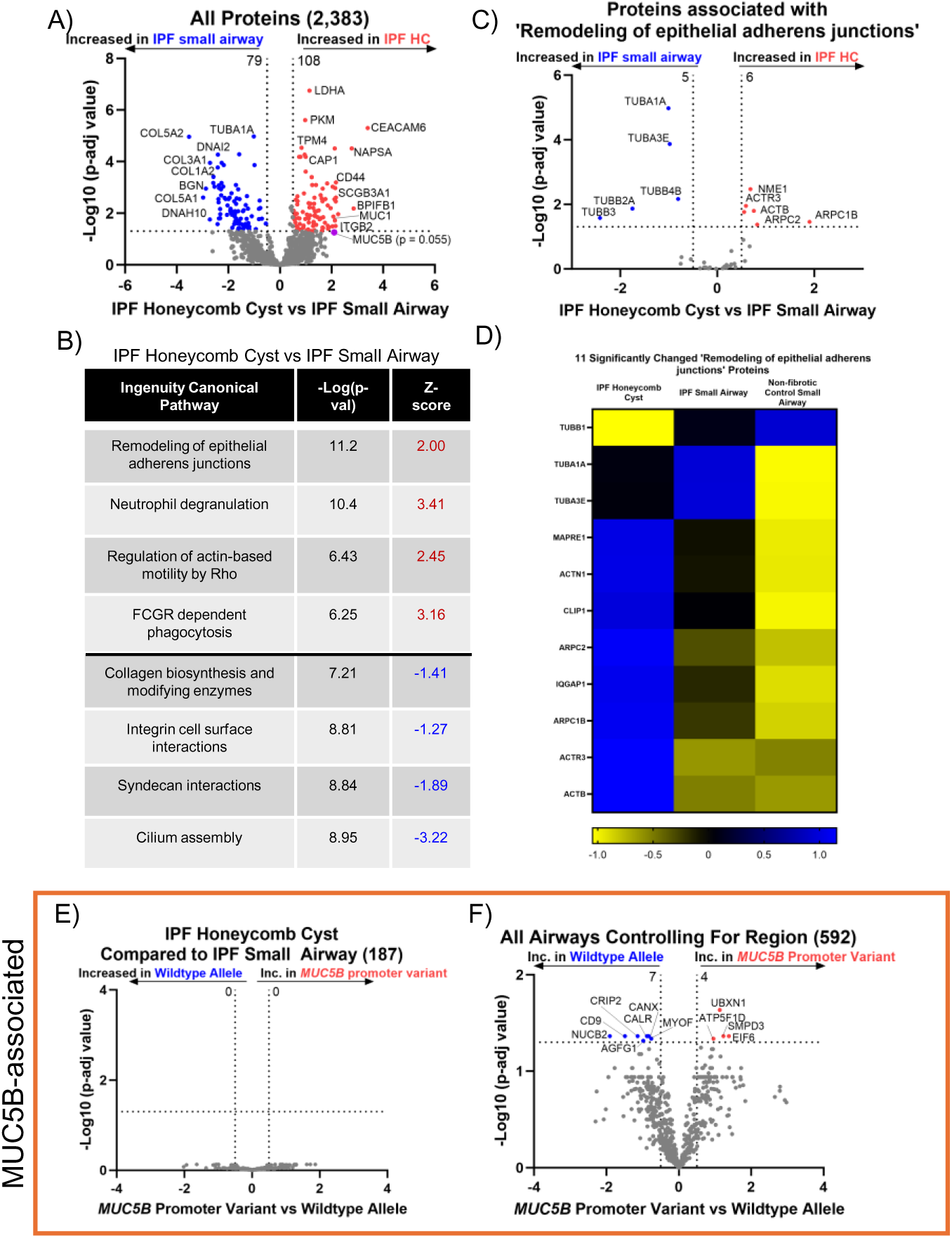
IPF honeycomb cysts are defined by remodeling of epithelial adherens junctions. (**A & C**) Volcano plot comparing IPF honeycomb cyst and IPF small airways, (C) and the subset of proteins associated with ‘remodeling of epithelial adherens junctions’, showing the negative natural log of p-adjusted values plotted against the base 2 log (fold change) for each protein. (**B**) Ingenuity pathway analysis showing the top 4 most upregulated (positive Z-score in red font) or top 4 most downregulated (negative Z-score in blue font) pathways. (**D**) A heatmap displaying Z-scores on the significantly changed ‘remodeling of epithelial adherens junctions’ proteins comparing IPF HC, IPF small airways, and non-fibrotic control small airways. (**E and F**) Volcano plots comparing *MUC5B* promoter variant and wildtype allele on (E) the subset of significantly changes IPF honeycomb cyst proteins in (A) and (F) the subset of all significantly changed airway proteins in IPF and non-fibrotic controls while controlling for region. N= 12 per group (6 wildtype and 6 *MUC5B* promoter variant).

We next focus on the proteins associated with ‘Remodeling of epithelial adherens junctions’ when comparing between IPF honeycomb cyst and IPF small airway (**Figure 4C**) and when comparing non-fibrotic control small airways to IPF airways (**Figure 4D**). Some notably increased proteins are associated with actin-related protein (ARP) 2/3 complex (ARPC1B, ARPC2, ARPC4, ACTR3). ARP2/3 complex is important in generating actin networks to allow for several cellular processes, including motility, membrane-trafficking, and endocytosis [19]. Actin networks are coupled to the ECM through their adhesive contacts, which have been associated with ECM remodeling. When comparing IPF honeycomb cyst to non-fibrotic control small airway, we find that the most significantly decreased pathways relate to ECM remodeling (collagen degradation, assembly of collagen fibrils and other multimeric structures, and extracellular matrix homeostasis) (**Supplemental Figure 3**). Further research into actin networks, airway cell adhesion, and ECM remodeling are warranted.

Interestingly, none of the 187 significantly changed IPF honeycomb cyst proteins were regulated by the *MUC5B* promoter variant (**Figure 4E**). We next re-assessed whether any of the 592 significantly altered airway proteins, from all airway sample types (non-fibrotic control small airway, IPF small airway, and IPF honeycomb cyst), are impacted by the *MUC5B* promoter variant while controlling for region, as previously performed using spatial transcriptomics data [20]. We find 11 significantly changed MUC5B-associated proteins (**Supplemental Figure 4 & Figure 4F**). CD9 and myoferlin (MYOF) are increased in IPF honeycomb cysts at the level of proteomics; we confirmed these findings using IHC (**Figure 5**). We next assessed CD9 and MYOF IHC expression in specimens with and without the *MUC5B* promoter variant in IPF specimens (**Supplemental Figure 5**). We found that the expression in IPF honeycomb cysts are highly variable, which may reflect a limitation of antibody-based approaches as compared to the high-sensitivity of mass spectrometry [21].

**Figure 5:**
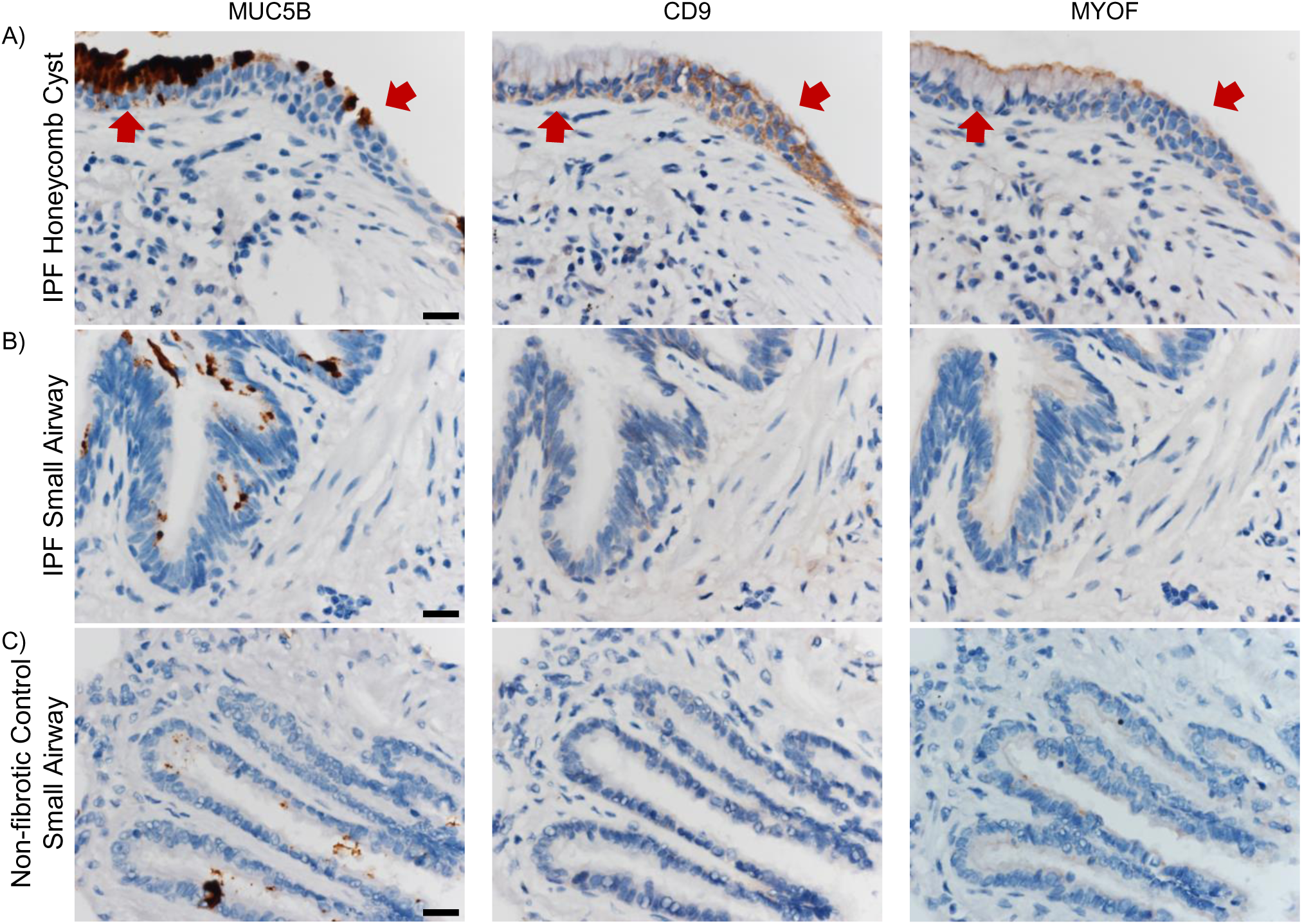
Airway expression of adhesion-associated proteins CD9 and myoferlin. Representative immunohistochemical stain for mucin 5B (MUC5B), CD9, and Myoferlin (MYOF) in (**A**) IPF honeycomb cyst (red arrows), (**B**) IPF small airway, and (**C**) control small airway. (Scale bar represents 20 microns)

### Fibroblastic foci (FF) are defined by increased translational control

We next sought to understand the proteomic signature of the IPF FF. A limitation to understanding the biological function of the FF is the lack of a fibroblastic structure in control lungs, therefore we compared the FF to adjacent alveolar structures which have been previously done [8, 22]. We identified 757 uniquely expressed proteins in IPF FF (**Supplemental Figure 2C**). Gene enrichment analysis demonstrated pathways involved in translational control (ribosomal scanning and start codon recognition, translation initiation complex, and activation of the mRNA upon binding of the cap-binding complex and eIFs) and ECM (collagen biosynthesis, collagen formation, and ECM proteoglycans) are enriched in the FF.

We next performed a quantitative analysis of the IPF FF as compared to adjacent IPF alveoli (**Figure 6A - B**). Increased proteins in the IPF FF are ECM-related, such as fibulin-2 (FBLN2), latent TGF-β binding protein 1 (LTBP1), and collagen XIV (COL14A1). The most significantly decreased protein is Na(+)/H(+) exchange regulatory co-factor NHE-RF2 (NHERF2), a protein primarily expressed in endothelial cells and not fibroblasts (IPF Cell Atlas). In accord with our qualitative analysis, pathway analysis of the quantitative data also demonstrates an increase in eukaryotic translation initiation. When comparing IPF FF to non-fibrotic control alveoli (**Supplemental Figure 6**), we additionally found increased translation control, including eukaryotic translation initiation, elongation, and termination. Among decreased pathways in IPF FF include negative regulators of translational control, such as PTEN signaling and mTOR regulation.

**Figure 6:**
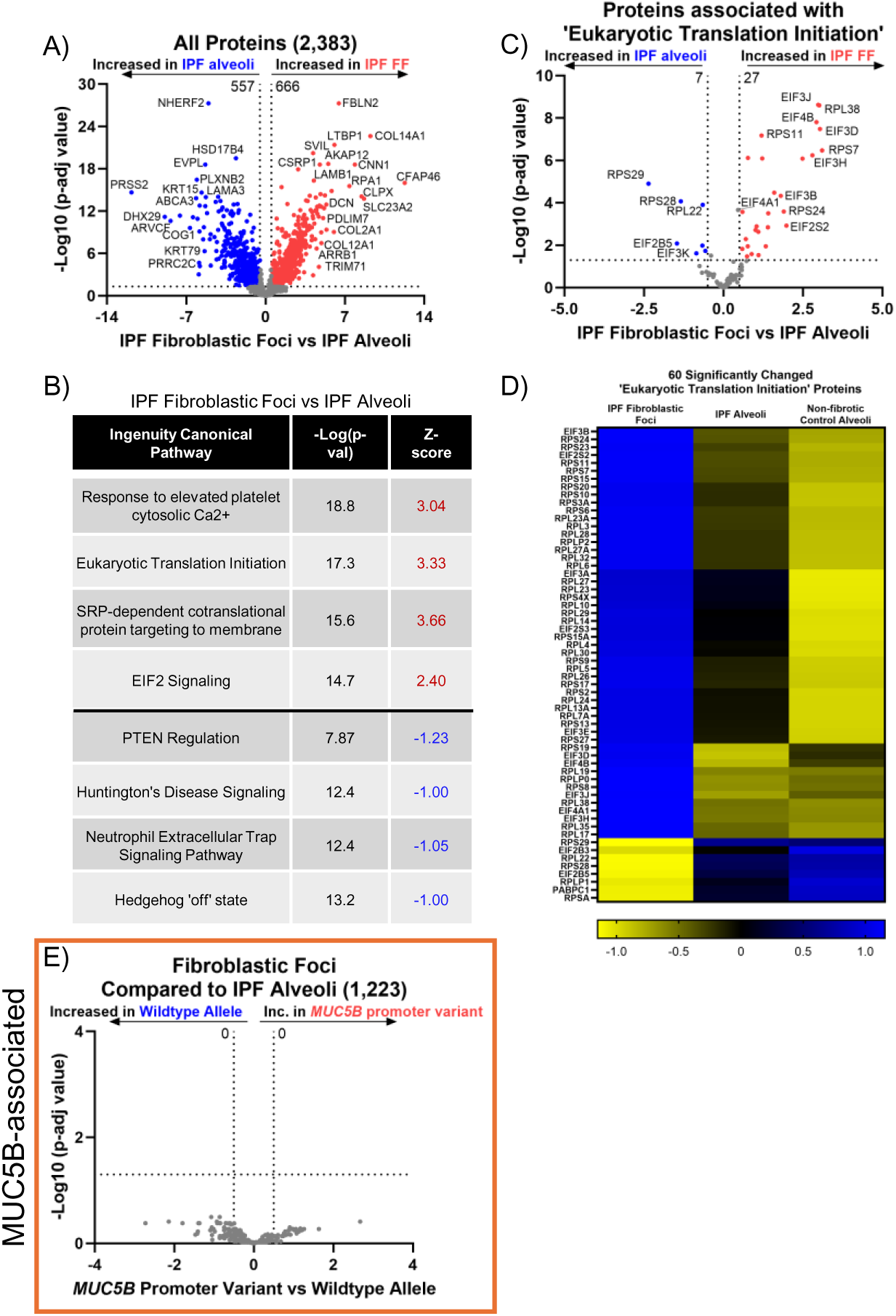
IPF fibroblastic foci are defined by increased translational control. (**A, C, & E**) Volcano plot comparing (A) IPF FF and IPF alveoli, (C) and the subset of proteins associated with ‘eukaryotic translation initiation’, showing the negative natural log of p-adjusted values plotted against the base 2 log (fold change) for each protein. (**B**) Ingenuity pathway analysis showing the top 4 most upregulated (positive Z-score in red font) or top 4 most downregulated (negative Z-score in blue font) pathways per comparison. (**D**) A heatmap displaying Z-scores of the significantly changed ‘eukaryotic translation initiation’ proteins comparing IPF FF, IPF alveoli, and non-fibrotic control alveoli. N = 12 per group (n = 20 IPF FF, balanced for the *MUC5B* promoter variant).

We next focus on the proteins comprising eukaryotic translation initiation (**Figure 6C – D**). The initiation of translation, particularly the binding of eukaryotic initiation factor 4F (eIF4F) complex to the 5’ mRNA cap, is a critical rate-limiting step in the process of translating mRNA into protein [23]. The eIF4F complex is composed of eIF4A, -4E, and - 4G, with eIF4A showing a significant increase (log2 of 0.46) in the IPF FF. Additionally, eIF4B is also elevated in the FF and is associated with eIF4A. Prior work has shown that IPF-derived fibroblasts exhibit deranged translational control, and that ECM transcripts are translationally activated by fibroblasts when interacting with pathological ECM [24, 25]. In our comparison, we find that the most increased pathway in IPF FF is ‘collagen biosynthesis and modifying enzymes’, with a Z-score of 4.26. Furthermore, we identified 119 ECM proteins that show significant changes in composition when comparing non-fibrotic control alveoli to IPF FF (**Supplemental Figure 6**). Thus, the proteomic signature of IPF FF is likely reflective of enhanced translational control that favors fibroblast translation of ECM transcripts. However, none of the 1,223 differentially expressed proteins associated with IPF FF are regulated by the *MUC5B* promoter variant (**Figure 6E**).

### Cellular stress defines the epithelia overlying the IPF fibroblastic foci

We next sought to define the IPF epithelia overlying FF and performed a gene enrichment analysis on the 383 uniquely expressed proteins (**Figure 7A**). We find that the most significantly increased pathway is ‘cellular response to stress’. This is consistent with a prior report showing that the epithelium lining the FF positively expresses endoplasmic reticulum (ER) stress mediator activation transcription factor 4 and 6 (ATF4 and ATF6, respectively) [26].

**Figure 7:**
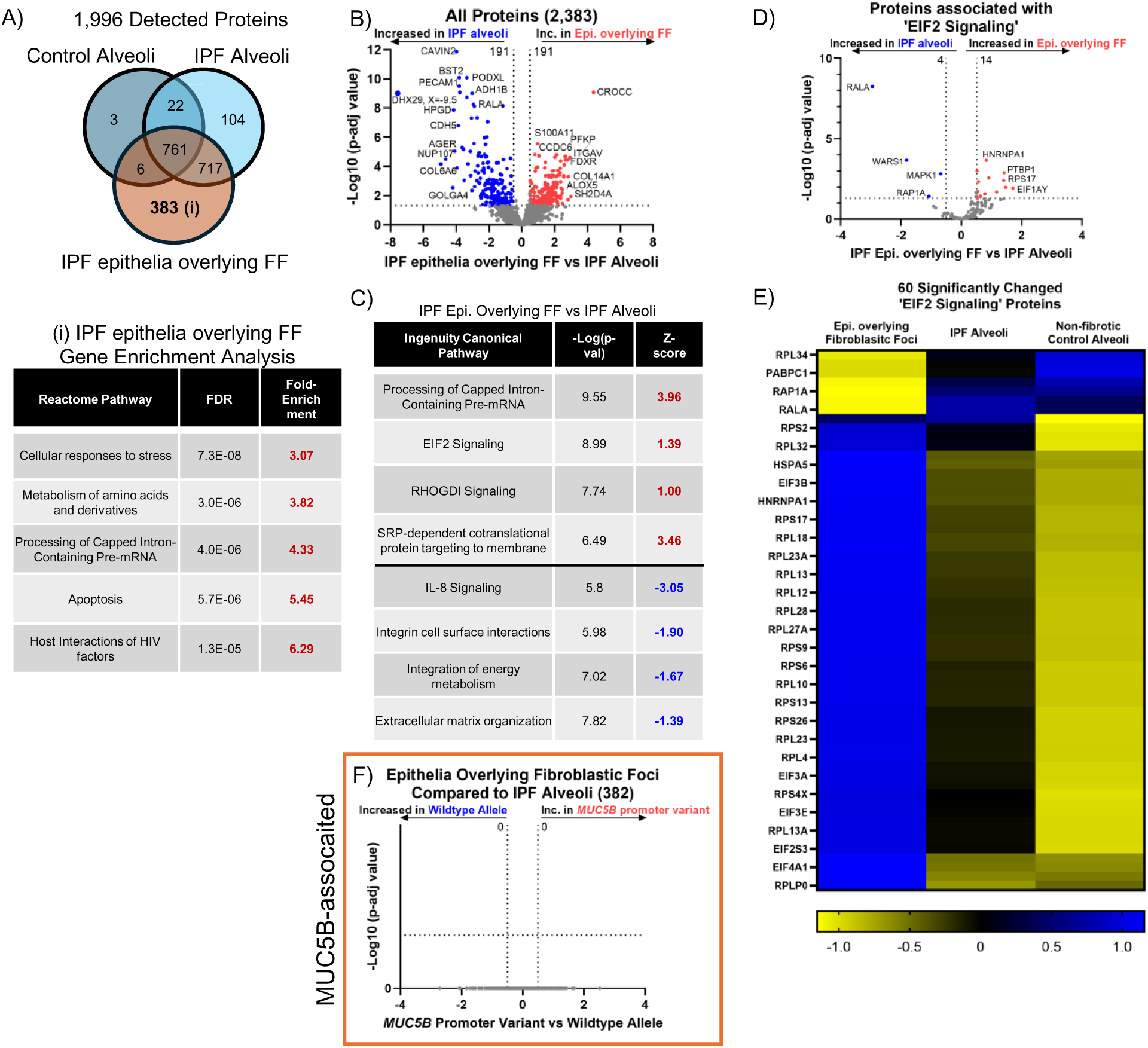
IPF epithelia overlying the FF is defined by cell stress pathways. (**A**) Venn diagram of detected proteins in the IPF epithelia overlying FF as compared to IPF and non-fibrotic control alveoli. Below is a gene enrichment analysis against the 383 uniquely expressed proteins in the epithelia overlying FF using www.pantherdb.org. (**B, D, & F**) Volcano plot comparing (B) IPF epithelia overlying the FF and IPF alveoli, (D) and the subset of proteins associated with ‘EIF2 signaling’, and (F) the subset of significantly changed proteins in (A) reassessed for the *MUC5B* promoter variant, showing the negative natural log of p-adjusted values plotted against the base 2 log (fold change) for each protein. (**C**) Ingenuity pathway analysis showing the top 4 most upregulated (positive Z-score in red font) or top 4 most downregulated (negative Z-score in blue font) pathways per comparison. (**E**) A heatmap displaying Z-scores of the significantly changed ‘EIF2 signaling’ proteins comparing IPF epithelia overlying FF, IPF alveoli, and non-fibrotic control alveoli. N = 12 per group (6 wildtype and 6 *MUC5B* promoter variant).

To further determine the functionality of the epithelia overlying the FF, we first compared these cells to IPF alveoli (**Figure 7B – C**). Pathway analysis demonstrated that ‘Processing of capped intron-containing pre-mRNA’ and ‘EIF2 Signaling’ are the most over-represented pathways in the epithelia overlying FF. Signaling through eIF2 has been shown to selectively translate mRNAs related to ER stress, such as ATF4 [27]. Our proteomic result support a recent spatial transcriptomic data showing that the transitional regions of IPF lungs, characterized by enrichment of FF, exhibit elevated ‘EIF2 signaling’ [28]. We next focused on the proteins associated with EIF2 signaling (**Figures 7D – E**). The most significantly increased protein in the epithelia overlying FF is heterogeneous nuclear ribonucleoprotein A1 (HNRNPA1), a protein that is sequestered to stress granules upon cellular stress [29]. Other stress granule proteins increased in the epithelia overlying the FF include eIF3A, eIF3B [30]. We further compared the epithelia overlying FF to non-fibrotic control alveoli (**Supplemental Figure 7**). We find that the most increased pathways relate to translational control. Together with increased ER stress and translational control, the epithelia overlying the FF appear to play a vital role in fibroblast activation.

To validate our spatial proteomic approach, we performed IHC on the IPF epithelia overlying the FF. We immunostained for cytokeratin 17 (marker of the epithelia overlying FF), phospho-eIF2alpha at Serine 51 and its downstream target ATF4 [31], and ATF6 (**Figure 8**). In accord with prior reports [26], we find positivity of these markers, whereas these stains were less prominent in adjacent IPF alveoli or non-fibrotic control alveoli. Similar to previous findings, none of the 382 proteins that exhibited significant changes in the epithelia overlying FF were influenced by the *MUC5B* promoter variant (**Figure 7F**).

**Figure 8:**
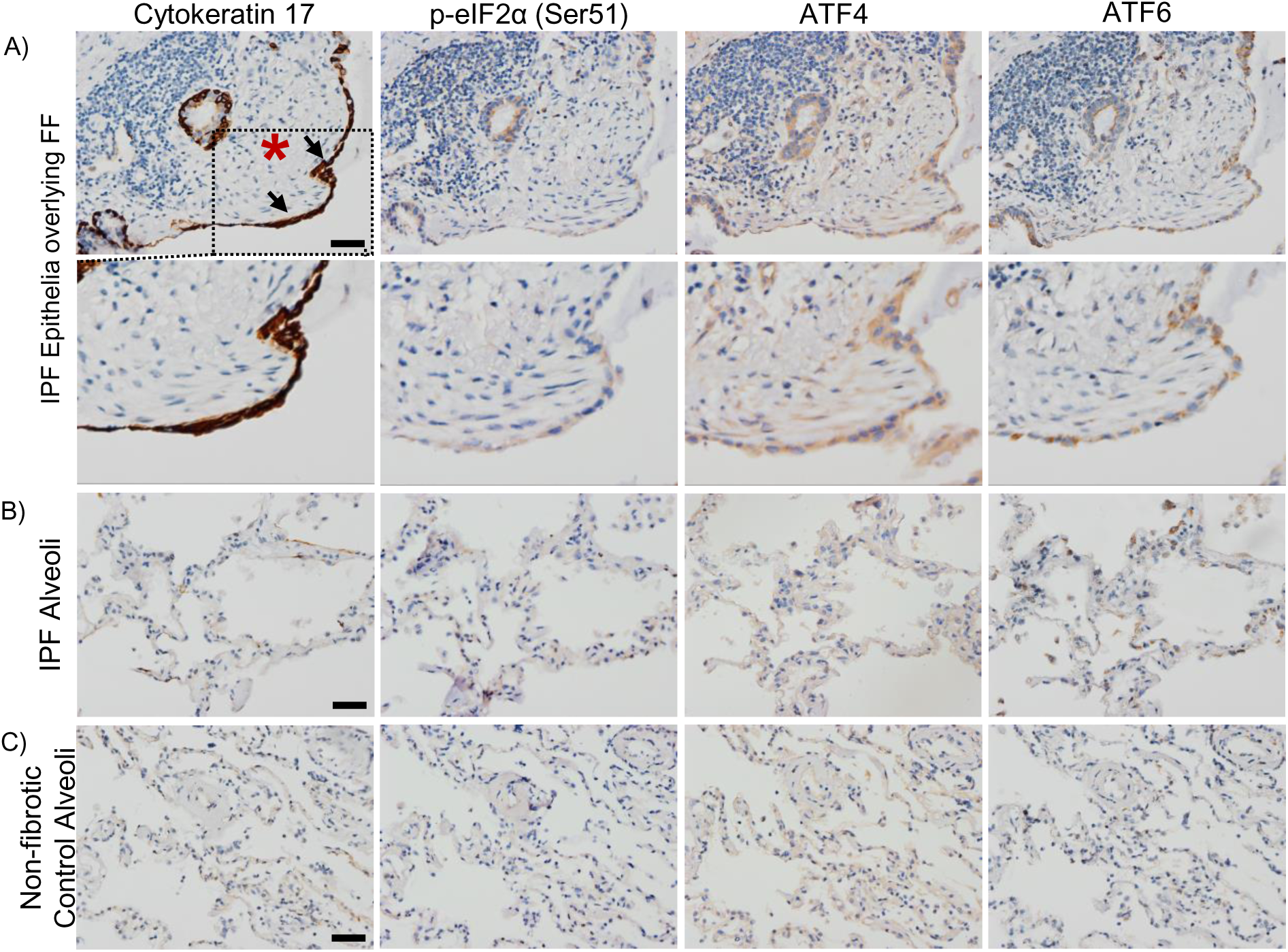
IPF epithelia overlying fibroblastic foci (FF) express stress markers. Representative images of immunohistochemical stain for cytokeratin 17, phospho-eIF2α (serine 51), ATF4, and ATF6 in (**A**) IPF epithelia (black arrows) overlying a fibroblastic foci (red asterisk) and (**B**) IPF alveoli, and (**C**) non-fibrotic control alveoli. Scale bar represents 50 microns.

## Discussion

Our primary goal was to determine whether the *MUC5B* promoter variant impacts the proteomic signatures of non-fibrotic control and IPF lungs. Our findings reveal that most proteomic changes associated with the *MUC5B* promoter variant are found in non-fibrotic control lungs, particularly in the alveoli. In contrast, we found little evidence linking the *MUC5B* promoter variant to proteomic alterations in IPF lungs. This is likely attributed to the dominant fibrotic signature masking the effect of the *MUC5B* promoter in the IPF lung and our relatively small sample size (n = 12 non-fibrotic control or n = 12 IPF specimens, with 6 harboring the *MUC5B* promoter variant). Given that the *MUC5B* promoter variant is the dominant risk factor for developing IPF, our findings suggest that enhanced expression of *MUC5B* in non-fibrotic lungs may induce early changes that predispose them to lung fibrosis by establishing a vulnerable bronchoalveolar epithelium. This is further supported by recent findings demonstrating that enhanced expression of Muc5b in either AT2 or airway cells alone does not drive fibrosis *in vivo*, at least in short-term bleomycin models of lung fibrosis. However, following bleomycin injury, Muc5b overexpression resulted in enhanced collagen deposition, honeycomb cyst formation and mucus production [32, 33].

Interestingly, IL3-signaling is increased in the non-fibrotic control alveoli in association with the *MUC5B* promoter variant. IL-3 is a cytokine produced by T-lymphocytes and mast cells that stimulate the development of a variety of immune cells and plays a major role in inflammation [34]. In experimental rodent models of acute lung injury, IL-3 knockout reduced proinflammatory mediators and neutrophil abundance [35]. Thus, enhanced expression of *MUC5B* in non-fibrotic lungs may involve early inflammatory changes.

The unfolded protein response (UPR) is a type of ER stress that occurs when protein processing is disturbed, leading to the accumulation of misfolded proteins. ER stress is a significant factor in lung fibrosis and has been previously associated with genetic risk variants (e.g. *SFTPC*) [36, 37]. Given that the gain-of-function *MUC5B* promoter variant has also been associated with ER stress in epithelial cells [32], *MUC5B* may contribute to this process. In fact, the most significantly increased MUC5B-associated protein in airways is UBX domain-containing protein 1 (UBXN1) (**Figure 4F**), a negative regulator of the UPR [38]. In addition, we find that the *MUC5B* promoter variant is associated with decreased expression of protein folding proteins calreticulin (CALR) and calnexin (CANX) in airway epithelium [39, 40]. A closer inspection of MUC5B-associated proteins in non-fibrotic control alveoli yielded several proteins associated with protein homeostasis (**Figure 1B**). For instance, BAG family molecular chaperone regulator 2 (BAG2) plays a prominent role in protein homeostasis by mediating protein refolding [41]. In addition, 26S proteasome regulatory subunit-8 (PSMC5) and subunit-7 (PSMC2) are involved in protein homeostasis by degrading misfolded proteins [42]. BAG2, PSMC5 and PSMC2 are MUC5B-assocated proteins that are decreased in non-fibrotic control alveoli. Thus, it is plausible that enhanced expression of *MUC5B* may lead to ER stress and UPR in lung epithelia that predisposes the lung to fibrosis.

Herein, we are the first to determine the proteomic signature of the epithelia overlying FF. We found that the epithelia overlying FF clustered with alveolar proteins and are the most distant to airway groups (**Figure 2**), suggesting that they share properties with alveolar cells. Secondly, gene enrichment analysis of the uniquely expressed IPF epithelia overlying FF proteins show ‘cell response to stress’ as the strongest category **(Figure 7A**) among other stress related pathways (e.g. eIF2 signaling) (**Figure 7C**). How the epithelium becomes stressed is unknown. We speculate that enhanced expression of *MUC5B* in lung epithelia may drive ER stress/UPR and the emergence of this cell population. Other groups have shown that AT2 cells, upon injury or the ablation of alveolar type I (AT1) cells, differentiated into a pre-alveolar type-1 transitional cell state (PATS) that share similarities with the epithelia overlying FF [43, 44]. Consequently, the accumulation of PATS is associated with activated alveolar fibroblast and ECM deposition. Given that fibroblastic foci are lined with ‘stressed’ epithelium, further work understanding *MUC5B*/ER stress/lung epithelium/mesenchymal crosstalk will likely enhance our understanding of fibroblastic foci development.

Our analysis revealed that IPF honeycomb cysts are defined by increased ‘remodeling of epithelial adherens junctions’, a pathway that was recently implicated in IPF distal airways using spatial transcriptomics [45]. Adherens junctions regulate cell polarity, extracellular matrix deposition, and collective cell migration [46, 47]. Collective cell migration is dysfunctional in IPF, resulting in increased migration compared to healthy counterparts [48, 49]; a phenotype that was recapitulated by lung injury *in vivo* [50]. Additionally, we find that CD9 adhesion protein is differentially regulated by the *MUC5B* promoter variant in airways (**Figure 4F**). CD9 is important for regulating epithelial collective cell migration [51]. Provided that ECM homeostasis is perturbed in IPF honeycomb cysts [7, 45] and that adherens junctions are influenced by ECM, this suggests a dynamic interplay between cell adhesion, adherens junctions, and ECM remodeling. Thus, fibrosis may be stimulated in an airway-centric manner following injury (e.g. smoking, *MUC5B*, etc), which modulates airway cell adhesions and triggers ECM remodeling.

## Conclusion

By utilizing LCM-MS, we found that the *MUC5B* promoter variant, the strongest risk factor for IPF development, predominantly impacts the proteomic profiles of non-fibrotic lung tissue. We propose that enhanced *MUC5B* expression in lung epithelial cells may prime the epithelium through ER stress/UPR pathways for a secondary injury to initiate fibrosis. Furthermore, we found that the epithelium overlying the IPF fibroblastic foci exhibited cellular stress pathways. These findings underscore the role of the *MUC5B* promoter variant in priming the lung for fibrosis and emphasize the need to target ER stress pathways to mitigate IPF progression.

## Materials and Methods

### Lung Tissue Procurement

All lung specimens met the criteria for IPF diagnosis following current guidelines [1]. Patients were consented and samples were approved for research by the Colorado Multiple Institutional Review Board (COMIRB # 15-1147) and through the Lung Tissue Research Consortium (LTRC). Fresh lung tissue was fixed in 4% formaldehyde for 48 hours and then transferred into 70% ethanol prior to processing into formalin-fixed paraffin-embedded (FFPE) blocks.

### Laser Capture Microdissection

We utilized an Olympus IX63 microscope integrated with Molecular Machinery Instruments (MMI) technologies to perform laser capture microdissection. FFPE human lung tissue was sectioned at 5 microns and collected onto Molecular Machines & Industries (MMI) membrane slides (catalog 50103) and stained with routine hematoxylin and eosin (H&E). Using MMIs caplift technology, we captured regions of interest into MMI tubes (MMI, 50204) and stored these samples in - 80°C prior to processing for mass spectrometry. Samples stayed in the freezer for a maximum of 3 months prior to processing. For all samples, we collected volumes of 0.005 – 0.010 mm^3^.

### Sample Processing for Mass Spectrometry

Laser captured material were processed as previously described [6–8] with minor modifications that we highlight here. To achieve sample shearing, we utilized Covaris S220 and Sonolab 7.2 software at a setting of peak power of 200, duty factor of 20.0, cycle bursts of 200, with a duration of 50 seconds followed by a 10 second cool period (water temperature set to 4 °C). This cycle was repeated 10 times for a total of 10 minutes. For sample digestion, we utilized 200 ng of trypsin within 25 uL of digestion buffer and incubated at 47°C for 2-hours. All other processing, digestion, and desalting steps were performed as previously described.

### Mass Spectrometry Acquisition

Desalted peptides were adjusted to a protein concentration of 50 ng/uL with 0.1% formic acid (FA) in preparation for MS analysis. Digested peptides were loaded into autosampler vials and analyzed directly using a NanoElute liquid chromatography system (Bruker, Germany) coupled with a timsTOF SCP mass spectrometer (Bruker, Germany). Peptides were separated on a 75 µm i.d. × 25 cm separation column packed with 1.6 µm C18 beads (IonOpticks) over a 90-minute elution gradient. Buffer A was 0.1% FA in water and buffer B was 0.1% FA in acetonitrile. Instrument control and data acquisition were performed using Compass Hystar (version 6.0) with the timsTOF SCP operating in parallel accumulation-serial fragmentation (PASEF) mode under the following settings: mass range 100-1700 m/z, 1/k/0 Start 0.7 V s cm-2 End 1.3 V s cm-2; ramp accumulation times were 166 ms; capillary voltage were 4500 V, dry gas 8.0 L min-1 and dry temp 200°C. The PASEF settings were: 5 MS/MS scans (total cycle time, 1.03 s); charge range 0–5; active exclusion for 0.2 min; scheduling target intensity 20,000; intensity threshold 500; collision-induced dissociation energy 10 eV.

### Data Processing

Data was searched using MSFragger via FragPipe v 20.0. Precursor tolerance was set to ±15 ppm and fragment tolerance was set to ±0.08 Da. Data was searched against SwissProt restricted to *Homo sapiens* with added common contaminants (20,410 total sequences). Enzyme cleavage was set to semi-specific trypsin for all samples. Fixed modifications were set as carbamidomethyl (C). Variable modifications were set as oxidation (M), oxidation (P) (hydroxyproline), Gln->pyro-Glu (N-term Q), and acetyl (Peptide N-term). Results were filtered to 1% FDR at the peptide and protein level. The data was then uploaded and processed with FragPipe-Analyst using LFQ settings, no normalization, p-value cutoff at 0.05, Perseus-type imputation and FDR correction using Benjamini Hochberg to acquire a dataset for further statistical processing.

### Statistical Methods

For the qualitative analysis, proteins were considered detected if it was detected in 80% or more of samples within each group. For quantitative analysis, we used a linear mixed model framework to test for differences between the *MUC5B* promoter variant or between region groups and account for repeated measures within subjects. Proteins were removed if they were undetected in greater than 24 samples. For the remaining proteins, we performed quantile normalization on the imputed matrix. To test for differential abundance in the quantitative analyses, we modeled the quantile normalized values by region group including a random effect for subject using the R package lmerSeq [52]. For the *MUC5B* promoter variant comparisons, data was subset to each region and tested for differentially abundant proteins between subjects with (GT) and without the risk variant (GG). We excluded results that had minimal variance (determined to be singular). Multiple testing correction was performed using Benjamini-Hochberg adjustment and statistical significance was set to an alpha level of 0.05.

### Immunohistochemistry

FFPE lung tissue was sectioned at 5 microns and de-paraffinized by submerging in a series of xylene and alcohol baths. Slides underwent antigen heat retrieval in either Universal HIER solution (Abcam; ab208572) or EDTA pH 8.0 for 20 minutes in a steamer and then allowed to cool for 20 minutes to room temperature. Slides are then treated with 3% hydrogen peroxide for 10 minutes and blocked for 1 hour (ThermoFisher; SuperBlock; 37535). Primary antibody was added overnight in 10% SuperBlock solution. Antibodies used: phosphor-EIF2alpha (CellSignal, #3398S; EDTA 1:50 dilution), Cytokeratin 17 (Abcam; ab53707; HIER 1:16,000 dilution), ATF4 (Proteintech; 10835-1-AP; Citrate 1:16,000 dilution), ATF6 (Abcam; ab227830; HIER 1:1,500 dilution), MUC5B (Novusl NBP2-50522; HIER 1:8,000), STAT5A (Abcam; AB106095; 1: 15,000), CD9 (Abcam; ab2215, HIER 1:80,000), and MYOF (Invitrogen; PA5-53134; HIER 1:8,000). On the next day, we used Novolink Polymer Kit (Leica Biosystems; RE7200-CE) following manufacturers’ recommendations. Finally, we reacted the slides with DAB substrate (Abcam; ab64238) for 5 minutes followed by hematoxylin counterstaining and coverslipping.

### Microscopy

Brightfield images were taken with an Olympus BX63 microscope using CellSens Dimensional Software.

### Data Availability

Raw mass spectrometry data will be uploaded onto the public proteomic repository ‘ProteomeXchange’ upon final publication.

## Author Contributions

J.A.H. and D.A.S. conceived and supervised the project. J.A.H., M.M., and R.B. performed LCM-MS. J.B. procured lung specimens and C.D.C. reviewed lung pathology. J.A.H., M.M., and R.Z.B. analyzed data. J.A.H. and N.Y.L. performed immunohistochemistry. J.A.H. wrote the manuscript with input from all the authors (M.M, R.Z.B., R.B., N.Y.L., J.B., C.D.C., J.P.H., J.S.K., C.M.M., K.C.H., I.V.Y., and D.A.S.).

