## Supplemental Figures for "The *MUC5B* promoter variant results in proteomic changes in the non-fibrotic lung"

### Supplemental Figure 1

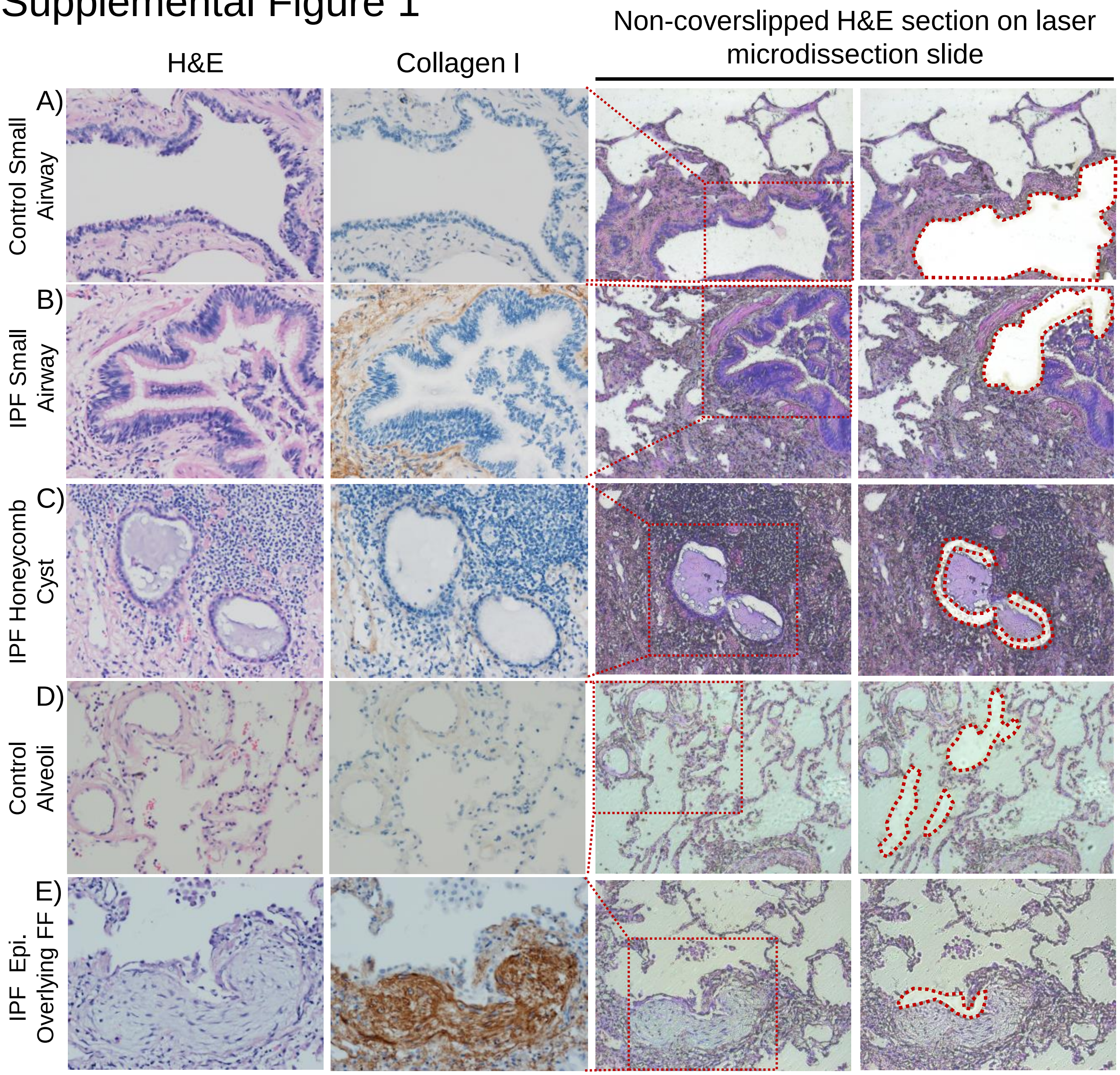


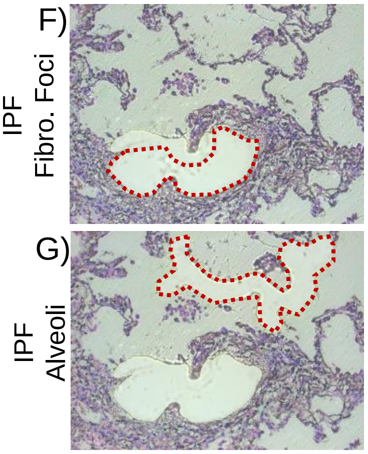


**Supplemental Figure 1: Our laser capture microdissection approach.** Herein we show our capacity to precisely capture regions of interest using both H&E and collagen I as a guide. A representative image of the capture of a (**A**) non-fibrotic control airway, (**B**) IPF small airway (**C**) IPF honeycomb cyst, and (**D**) non-fibrotic control alveoli. In the next set of panels, we laser capture the (**E**) epithelia overlying fibroblastic foci (FF), (**F**) IPF FF, and (**G**) IPF alveoli from the same region. The furthest right panel shows the regions captured (outlined by a red dotted line).

### Supplemental Figure 2

A)

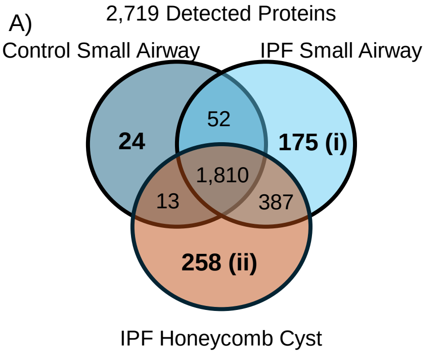


**(i)** IPF Small Airway
Gene Enrichment Analysis

| Reactome Pathway | FDR | Fold-Enrichment |
|---|---|---|
| Anchoring of the basal body to the plasma membrane | 7.8E-05 | 11.85 |
| Cilium Assembly | 4.7E-04 | 6.83 |

**(ii)** IPF Honeycomb Cysts
Gene Enrichment Analysis

| Reactome Pathway | FDR | Fold-Enrichment |
|---|---|---|
| Membrane Trafficking | 1.7E-03 | 3.06 |
| Neutrophil degranulation | 3.0E-03 | 3.2 |
| Innate Immune System | 1.2E-02 | 2.2 |
| RHOQ GTPase cycle | 4.9E-02 | 7.78 |

B)

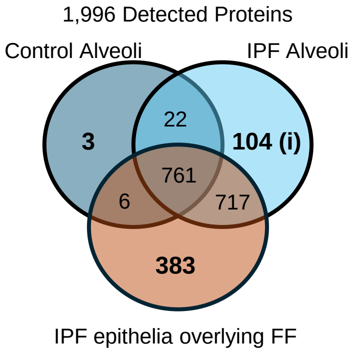


**(i)** IPF Alveoli
Gene Enrichment Analysis

| Reactome Pathway | FDR | Fold-Enrichment |
|---|---|---|
| Regulation of insulin secretion | 3.3E-05 | 20.11 |
| Integration of energy metabolism | 1.1E-04 | 14.66 |
| G-protein activation | 5.0E-03 | 32.69 |
| Thromboxane signalling through TP receptor | 6.0E-03 | 32.69 |
| ADP signaling through P2Y purinoceptor 1 | 7.5E-03 | 32.69 |

C)

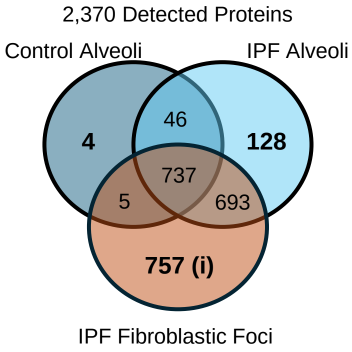


**(i)** IPF Fibroblastic Foci
Gene Enrichment Analysis

| Reactome Pathway | FDR | Fold-Enrichment |
|---|---|---|
| Non-integrin membrane-ECM interactions | 4.6E-07 | 6.67 |
| Ribosomal scanning and start codon recognition | 3.3E-06 | 6.22 |
| Translation initiation complex formation | 1.7E-05 | 5.78 |
| Activation of the mRNA upon binding of the cap-binding complex and eIFs | 1.9E-05 | 5.68 |
| Collagen biosynthesis and modifying enzymbes | 1.2E-05 | 5.48 |
| Signaling by MET | 1.0E-05 | 5.17 |
| ECM Proteoglycan | 1.1E-05 | 5.17 |
| Collagen formation | 3.5E-06 | 5.01 |

**Supplemental Figure 2: Qualitative spatial analysis of IPF.** Venn diagrams of detected proteins in (**A**) airway (**B**) alveolar, and (**C**) fibroblastic foci. A protein was considered detected if found in 80% of samples per group (left diagrams). A gene enrichment analysis of the uniquely expressed proteins using www.pantherdb.org is shown on the right. For the fibroblastic foci, we focused on the 50 most significantly regulated pathways of 357 arranged by fold-enrichment. N = 12 per group (n = 20 for fibroblastic foci).

### Supplemental Figure 3

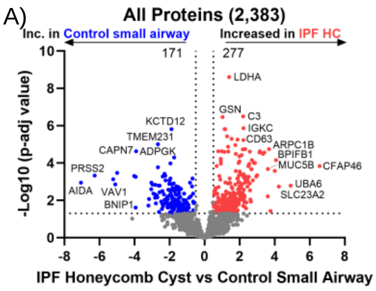


B) IPF Honeycomb Cyst vs Control Small Airway

| Ingenuity Canonical Pathway | -Log(p-val) | Z-score |
|---|---|---|
| Neutrophil degranulation | 24.0 | 4.16 |
| Response to elevated platelet cytosolic Ca2+ | 16.4 | 4.49 |
| Eukaryotic Translation Elongation | 16.2 | 4.15 |
| SRP-dependent cotranslational protein targeting to membrane | 15.5 | 4.26 |
| Extracellular matrix organization | 4.5 | -1.90 |
| GP6 Signaling Pathway | 4.54 | -1.90 |
| Assembly of collagen fibrils and other multimeric structures | 4.75 | -1.41 |
| Collagen degradation | 5.71 | -1.67 |

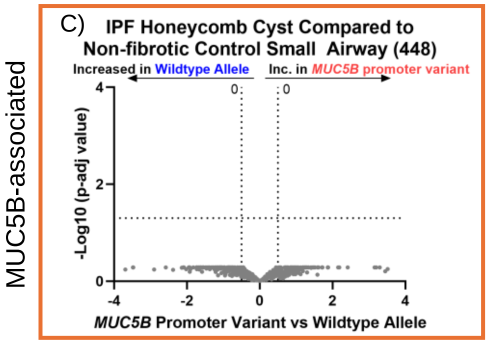


**Supplemental Figure 3: IPF honeycomb cysts demonstrate deranged ECM remodeling.** (**A & C**) Volcano plot comparing IPF honeycomb cysts and non-fibrotic control small airways (A) for all proteins and (C) the subset of significantly changed proteins from (A) comparing for the *MUC5B* promoter variant, showing the negative natural log of p-adjusted values plotted against the base 2 log (fold change) for each protein. (**B**) Ingenuity pathway analysis showing the top 4 most upregulated (positive Z-score in red font) or top 4 most downregulated (negative Z-score in blue font) pathways per comparison. N = 12 per group.

### Supplemental Figure 4

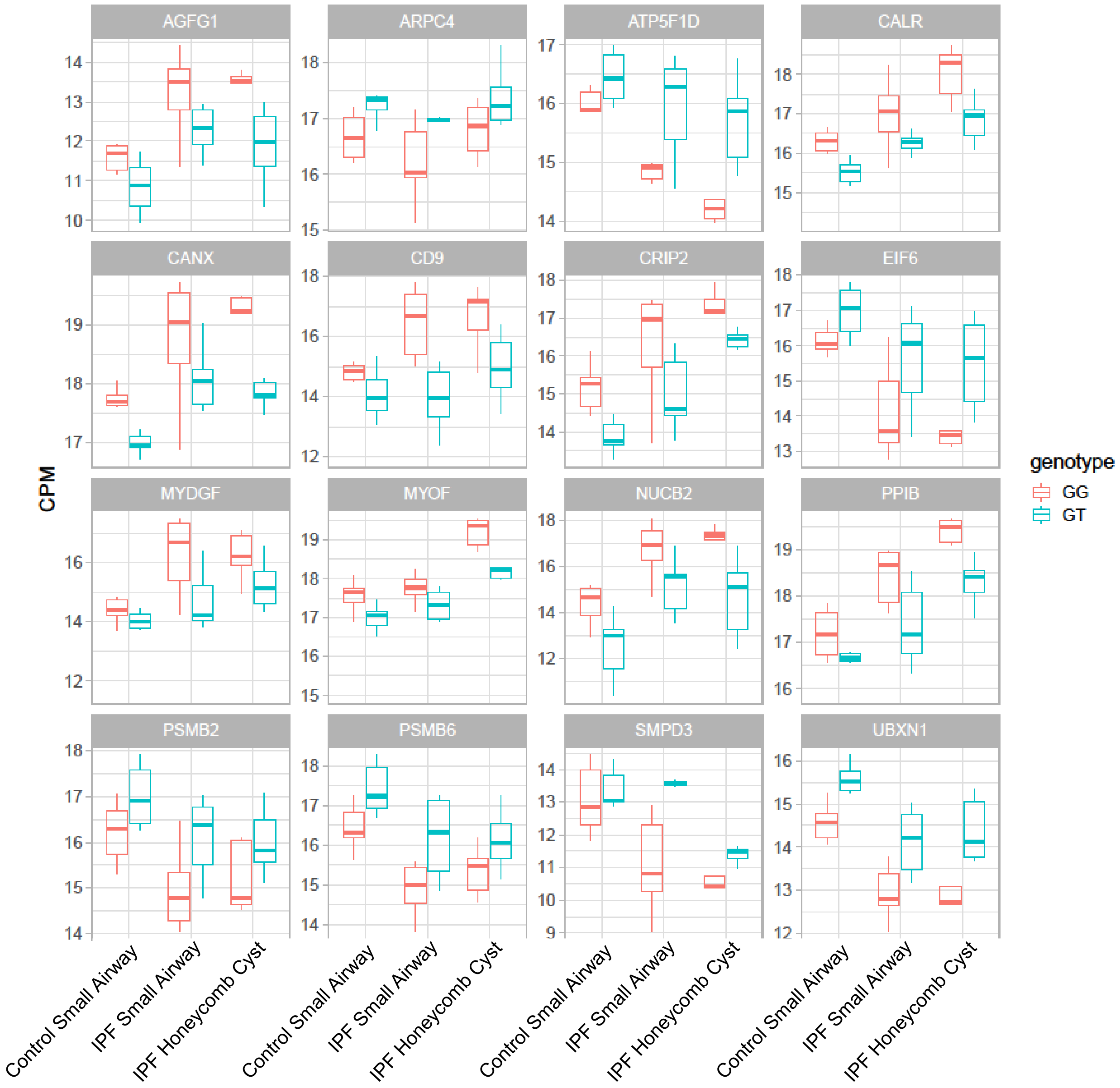


**Supplemental Figure 4: Impact of *MUC5B* promoter variant on the proteomic profiles of airways.** Shown is the affect of *MUC5B* promoter variant (GT) on control small airway, IPF small airway, or IPF honeycomb cyst (n = 6 per group). Although underpowered per group individually, our combined analysis when controlling for disease identified 11 significantly changed proteins when comparing for *MUC5B* promoter variant (original Figure 4F).

### Supplemental Figure 5

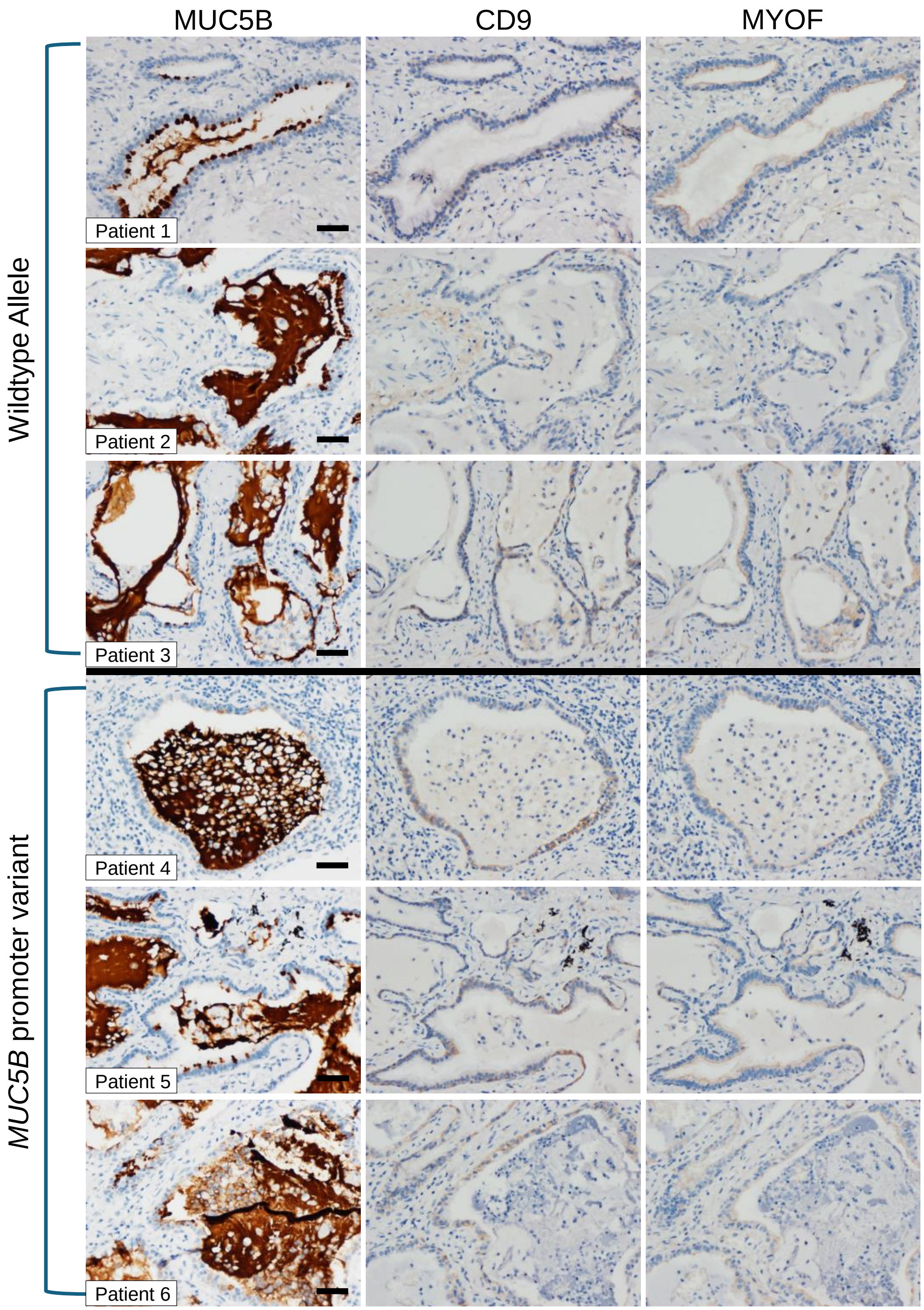


**Supplemental Figure 5: Expression of CD9 and MYOF in IPF honeycomb cyst:** Immunohistochemistry against MUC5B, CD9, and MYOF in 6 IPF specimens (3 wildtype allele and 3 *MUC5B* promoter variant). Scale bar represents 50 microns.

### Supplemental Figure 6

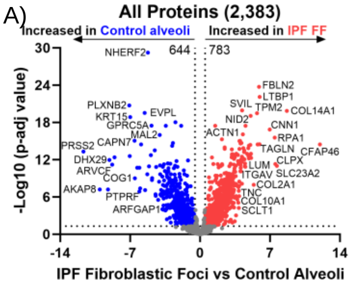


B)

IPF Fibroblastic Foci vs Control Alveoli

| Ingenuity Canonical Pathway | -Log(p-val) | Z-score |
|---|---|---|
| Eukaryotic Translation Initiation | 41 | 5.76 |
| SRP-dependent cotranslational protein targeting to membrane | 35.7 | 6.07 |
| Eukaryotic Translation Elongation | 35.1 | 5.37 |
| Eukaryotic Translation Termination | 34.2 | 5.286 |
| Processing of Capped Intron-Containing Pre-mRNA | 9.46 | -1.02 |
| mTOR Signaling | 11.7 | -1.39 |
| Mitochondrial dysfunction | 13.8 | -2.32 |
| Huntington's Disease Signaling | 14 | -2.00 |

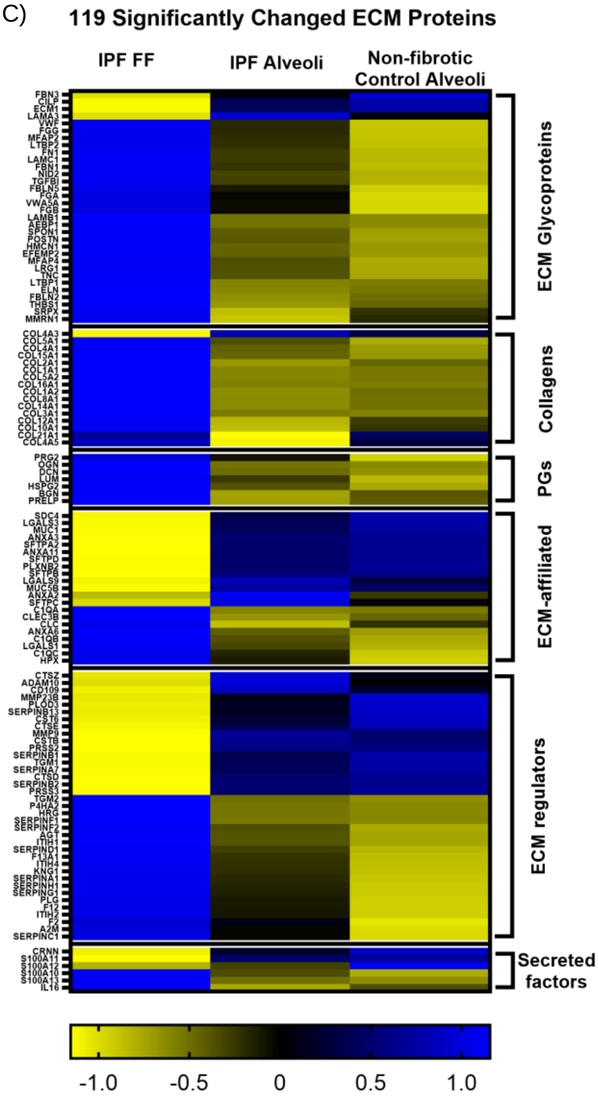


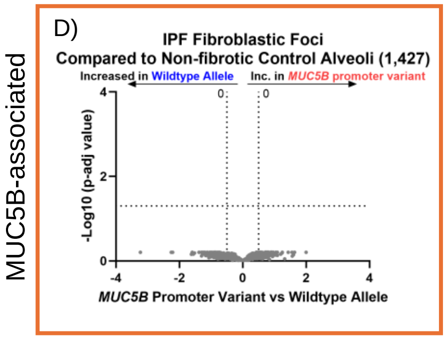


**Supplemental Figure 6: IPF fibroblastic foci are defined by increased translational control.** (**A & D**) (A)Volcano plot comparing IPF FF and non-fibrotic control alveoli and, (D) the subset of significantly changed proteins from (A) when comparing the *MUC5B* promoter variant, showing the negative natural log of p-adjusted values plotted against the base 2 log (fold change) for each protein. (**B**) Ingenuity pathway analysis showing the top 4 most upregulated (positive Z-score in red font) or top 4 most downregulated (negative Z-score in blue font) pathways. (**C**) A heatmap displaying Z-scores on the significantly changed extracellular matrix proteins categorized by their groups comparing IPF FF, IPF alveoli, and non-fibrotic control alveoli. N= 12 per group (n = 20 FF, balanced for the *MUC5B* promoter variant).

### Supplemental Figure 7

A)

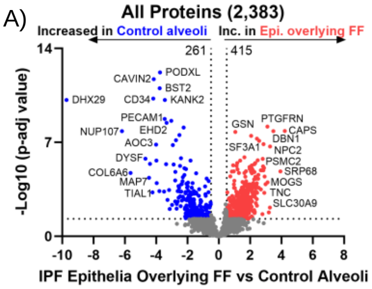


B)

IPF Epi. Overlying FF vs Control Alveoli

| Ingenuity Canonical Pathway | -Log(p-val) | Z-score |
|---|---|---|
| Eukaryotic Translation Initiation | 45.2 | 6.30 |
| SRP-dependent cotranslational protein targeting to membrane | 41.2 | 6.27 |
| Eukaryotic Translation Elongation | 40 | 6.25 |
| Neutrophil degranulation | 39.1 | 3.09 |
| Coronavirus Pathogenesis Pathway | 8.89 | -3.00 |
| Gene and protein expression by JAK-STAT signaling after IL-12 stimulation | 9.53 | -1.16 |
| Regulation of eIF4 and p70S6K Signaling | 13.8 | -1.00 |
| mTOR Signaling | 14.4 | -2.11 |

MUC5B-associated

C)

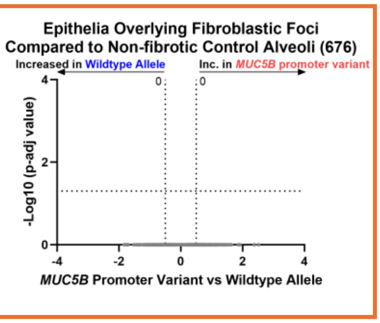


**Supplemental Figure 7: IPF epithelia overlying fibroblastic foci demonstrate increased translational control.** (**A & C**) Volcano plot comparing (A) IPF epithelia overlying FF and non-fibrotic control alveoli, and (C) the subset of significantly changed proteins in (A) assessed for the *MUC5B* promoter variant, showing the negative natural log of p-adjusted values plotted against the base 2 log (fold change) for each protein. (**B**) Ingenuity pathway analysis showing the top 4 most upregulated (positive Z-score in red font) or top 4 most downregulated (negative Z-score in blue font) pathways per comparison. N = 12 per group.
